# ACE: A Probabilistic Model for Characterizing Gene-Level Essentiality in CRISPR Screens

**DOI:** 10.1101/868919

**Authors:** Elizabeth R. Hutton, Christopher R. Vakoc, Adam Siepel

## Abstract

High-throughput knockout screens based on CRISPR-Cas9 are widely used to evaluate the essentiality of genes across a range of cell types. Here we introduce a probabilistic modeling framework, Analysis of CRISPR-based Essentiality (ACE), that enables new statistical tests for essentiality based on the raw sequence read counts from such screens. ACE estimates the essentiality of each gene using a flexible likelihood framework that accounts for multiple sources of variation in the CRISPR-Cas9 experimental process. In addition, the method can identify genes that differ in their degree of essentiality across samples using a likelihood ratio test. We show using simulations that ACE is competitive with the best available methods in predicting essentiality, and is especially useful for the identification of differential essentiality. Furthermore, by applying ACE to publicly available CRISPR-screen data, we are able to identify both known and previously overlooked candidates for genotype-specific essentiality, including RNA m**^6^**-A methyltransferases that exhibit enhanced essentiality in the presence of inactivating ***TP53*** mutations. In summary, ACE provides improved quantification of essentiality specific to cancer subtypes, and a robust probabilistic framework for identifying genes responsive to therapeutic targeting.

---

Pathogenic mutations and genotype-specific cancer liabilities can now be tested at unprecedented scales, owing to CRISPR-Cas9 (Clustered Regularly Interspaced Short Palindromic Repeats — CRISPR-associated protein 9) technology and large panels of diverse cell lines (1–10). These technologies allow genetic vulnerabilities to be demonstrated through direct perturbation, rather than through indirect associations from population enrichment studies or gene expression levels. Furthermore, in comparison to previous screens based on RNA interference, CRISPR knockout screens offer substantially improved sensitivity and specificity, due to increased effectiveness at disrupting targeted elements and reduced off-target effects (11–13).

In a typical CRISPR-Cas9 screen (Fig. 1a), the artificially expressed Cas9 nuclease introduces a double-stranded break in native DNA, guided by homology with the separately infected single-strand guide RNA (sgRNA). The cell then repairs the double-stranded break, typically inserting or deleting several nucleotides at the cut site in the process. The resulting genetic lesion often disrupts the function of the targeted region. The genetically modified cell is allowed to proliferate for a series of divisions, after which the effects on cell growth are measured by sequencing the sgRNA present in the final surviving population of cells. Lethal sgRNA constructs that have disrupted sites critical for normal cell growth will be underrepresented in the final population in contrast to ‘nonessential’ control sgRNAs. Moreover, differential essentiality can be detected from different levels of sgRNA depletion between samples. Hundreds of different cell lines have now been assayed in publicly available CRISPR knockout screens, offering an excellent opportunity to compare gene essentiality between panels of cancer cell lines according to their oncogenic mutations.

**Fig. 1.**
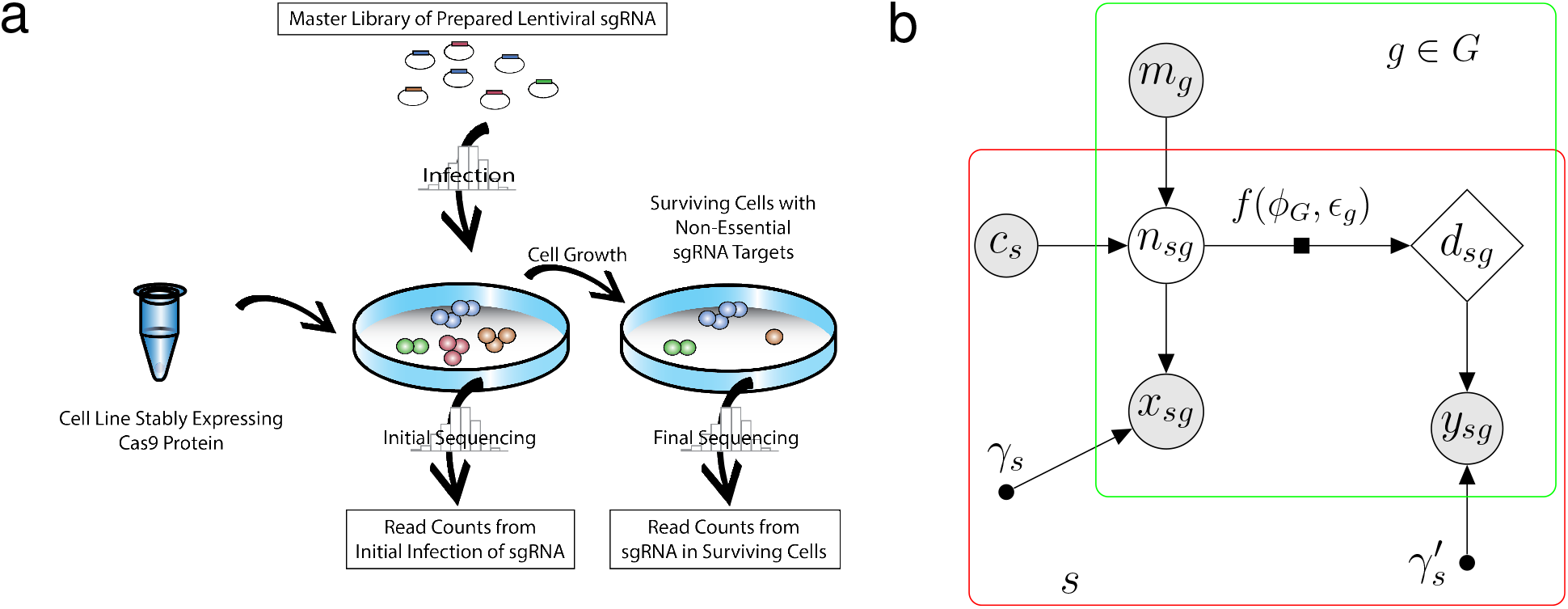
CRISPR-Cas9 Negative Selection Screen and Probabilistic Graphical Model. (a) The initial pool of Cas9-expressing cells is infected with a lentiviral sgRNA master library. The initial sgRNA abundances are measured by sequencing either the newly infected population of cells, the master library, or both. Cells are allowed to grow, and final sgRNA abundances are obtained by sequencing the surviving cells. (b) Graphical model illustrating how information is pooled across samples (*s*) and sgRNAs targeting each gene (*g* ∈ *G*) to infer each gene’s essentiality, *ϕ_G_*. From top-left to bottom-right, *m_g_* represents the fraction of the master library consisting of sgRNA *g*; *c_s_* epresents the number of cells in sample *s* (estimated by the user); *n_sg_* represents the number of cells initially infected with sgRNA *g* (such that *n_sg_ ≤ c_s_*); *d_sg_* represents the corresponding number of surviving cells after cell growth; and *x_sg_* and *y_sg_* represent the read counts obtained by sequencing *g* in the initial and surviving cells, respectively. The numbers of infected cells prior to (*n_sg_*) and following (*d_sg_*) growth are assumed to be related by the deterministic function *d_sg_* = *f* (*n_sg_*; *ϕ_G_, ϵ_g_*) = *n_sg_* (1 *− ϵ_g_ ϕ_G_*) (indicated by factor-graph notation). The efficiency *ϵ_g_* is determined by logistic regression and the gene essentiality values *ϕ_G_* are estimated by maximum likelihood (see Methods). The prior distribution for *n_sg_* and the sampling distributions for *x_sg_* and *y_sg_* are assumed to be Poisson. The scaling factors *γ_s_* and 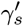 accommodate global properties of sequencing depth and cell growth, and are estimated in pre-processing (see Methods). Shaded nodes represent variables observed in the data, unfilled nodes represent latent variables, and smaller solid circles represent free parameters.

Within the general paradigm of a CRISPR-Cas9 screen, several variations of experimental design are possible. For example, many experiments evaluate the depletion of cells infected with lethal sgRNA by sequencing both the initial infected and the final population of cells (14–17). In contrast, other experiments streamline the process by sequencing only the initial pool of sgRNA constructs prior to infection (9, 18, 19), and use the sgRNA abundances in this ‘master library’ as a proxy for the initial frequencies. However, the sgRNA abundances in the master library vary by several orders of magnitude, and each biological replicate represents a separate infection into a new pool of cells, making this proxy imperfect. Overall, the variation in experimental design necessitates a flexible analytical framework that can robustly account for different sources of experimental error.

Through deliberate selection of the cell lines used, a CRISPR-Cas9 screen can reveal genes that are essential only in a specific oncogenic context, for example, in the presence of a loss-of-function mutation of an important tumor suppressor such as *TP53*. These context-dependent essential genes in cancer — sometimes called ‘non-oncogene addictions’ — are of particular interest because they may offer druggable therapeutic targets even when irreversible loss-of-function mutations to oncogenes are present (20). A better understanding of genotype-specific gene essentiality has the potential to enable patient-specific therapeutics that are tailored to mutations present at the time of diagnosis. Non-oncogene addiction has recently achieved prominence, for example, in the first round of chemical inhibitors for RNA epigenetic modifiers soon to enter phase I trials (21).

However, statistical tests for identifying such context-dependent essentiality require special care. Not only are they subject to reduced power because they require contrasting one subset of the data with another, but they can yield spurious results if the ‘test’ and ‘control’ subsets are not adequately matched in other respects. Thus, rigorous testing frameworks are needed that efficiently make use of the data and control for such biases.

Many of the statistical tests that have been applied to CRISPR-Cas9 screens rely on summary statistics such as average read counts across samples, or log-fold changes of read counts between the initial and final populations of cells. However, these summary statistics are incomplete descriptions of the raw data, leading to inevitable loss of power. In addition, these tests can be difficult to adapt to alternative experimental designs. For example, methods such as BAGEL and JACKS rely upon the log-fold change of read abundances (1, 2, 8–10, 22), but these log-fold changes may be calculated with respect to read counts obtained from either the initial infected population of cells or the master library, with potentially substantial impacts on the power of the tests. Another issue is that some methods reduce the estimation of gene essentiality to a binary classification of ‘essential’ or ‘nonessential’ genes (2, 4, 23), enabling the identification of clearly essential genes but blurring the comparison of weak and strong effects. Finally, most methods do not test directly for differential essentiality within sample subtypes (see Supplementary Fig. S1 for a comparison of methods).

In this paper, we introduce the Analysis of CRISPR-based Essentiality (ACE), a flexible probabilistic method that directly estimates gene-level essentiality values by maximum likelihood, accommodates variable experimental designs, and enables rigorous likelihood-based tests of both absolute and differential essentiality. The ACE model describes each experimental phase of the CRISPR screen — the master library infection, the initial and final sequencing — as a separate probabilistic process forming a hierarchical Bayesian model. The modularity of our framework permits the analysis of a variety of CRISPR-screen data sets while accounting for differences in experimental design. We validate ACE using both simulations and published sets of essential and nonessential genes. We then apply the method to publicly available data from Achilles DepMap (9, 24) and demonstrate several compelling examples of differential essentiality in non-small cell lung cancer, including both known and novel cases.

## Results

### Model Overview

ACE makes use of a hierarchical Bayesian model design to integrate information at the sample, gene, and individual sgRNA levels, with separate parameters at each level (see Fig. 1). The model is structured to account separately for the major sources of uncertainty at each stage of the experimental process. In addition, if samples are partitioned into ‘test’ and ‘control’ groups, ACE can estimate gene essentiality within each sample group separately, allowing for the detection of differential essentiality (discussed further below).

As illustrated in Fig. 1b, the structure of the ACE model reflects the experimental design of a CRISPR-Cas9 screen. The initial number of infected cells *n_sg_*, for each sgRNA *g* and sample (replicate) *s*, is assumed to be Poisson-distributed with an expected value given by the relative frequency of guide *g* in the master library, *m_g_*, scaled by the number of cells, *c_s_*, infected in the assay. Estimates of both *m_g_* and *c_s_* are provided by the user. The final number of infected cells, *d_sg_*, is then assumed to be given by a scaled version of the initial number, *d_sg_* = *n_sg_*(1 *− E_g_ϕ_G_*), where *ϵ_g_* ∈ [0,1] denotes the editing efficiency of sgRNA *g* and *ϕ_G_ ≤* 1 denotes the essentiality of gene *G*. These initial and final numbers of infected cells are not observed directly (they are latent variables) but are indirectly sampled via the sequencing process, resulting in observed read counts of *x_sg_* and *y_sg_*, respectively, which are provided by the user as inputs to the inference procedure. We assume Poisson-distributed read counts *x_sg_* and *y_sg_*, and sum over possible values of the latent variable *n_sg_* in the likelihood calculation (see Methods). Notably, the combination of the Poisson prior and the Poisson sampling distribution adequately captures the observed overdispersion in the read count data (see Supplementary Fig. S2). Independence is assumed across samples and replicates. The free parameters of the model are estimated by numerical maximization of the likelihood function (see Methods). The ACE software reports estimates of not only the essentiality of each gene, *ϕ_G_*, but also the efficiency of each sgRNA, *ϵ_g_*.

A value of *ϕ_G_* = 1 represents complete essentiality (driving *d_sg_* = *n_sg_*(1 *− ϵ_g_ϕ_G_*) to zero in the case of efficient editing, *ϵ_g_* = 1), whereas a value of *ϕ_G_* = 0 represents complete nonessentiality (allowing *d_sg_* = *n_sg_* even when *ϵ_g_ >* 0). Notably, however, our definition of *ϕ_G_* also allows for negative values, reflecting increased cell growth after inactivation of gene *G*. Such an increase might occur, for example, if a tumor suppressor gene were deactivated by guide RNAs. In practice, negative estimates of *ϕ_G_* are fairly frequently observed, either by random chance or from true increases in cell growth.

To pool information across sgRNAs and ensure identifiability of *ϵ_g_* and *ϕ_G_*, we determine *ϵ_g_* by logistic regression based on a user-defined set of features, and treat the regression coefficients only as free parameters (Methods). The efficiency of a class of sgRNA has been shown to be strongly correlated with a number of genomic features, including local GC content, proximity to heterochromatin, number of base mismatches, and abundance of off-target sites (18, 25–28). However, for simplicity and efficiency, we have used only the GC content of the sgRNA (one of the strongest predictors of efficiency) in our initial implementation. Nevertheless, ACE can easily be extended to consider additional features.

The two sample-specific scaling factors, *γ_s_* and 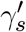, determine the relationships between the numbers of infected cells and the expected sgRNA counts in the initial and final data sets, respectively. These parameters apply globally to all sgRNAs and all genes. Rather than including these parameters in the numerical optimization of the likelihood function, we find it adequate to estimate them in preprocessing based on a median-of-ratios normalization. Notably, *γ_s_* is primarily a reflection of sequencing read depth, whereas 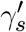is determined by both read depth and sample-specific factors in cell growth (see Methods). In the case where the bulk distribution is strongly influenced by essential genes, as when a library has a large number of essential targets, 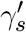 can alternatively be estimated from designated negative controls.

### Differential Essentiality

In the case of differential essentiality, ACE compares two (log) likelihoods for each gene *G*: a likelihood maximized under the constraint that the essentiality, *ϕ_G_*, is the same in designated ‘test’ and ‘control’ sample categories, and a likelihood that allows for different values of *ϕ_G_* in these two categories. These sample categories can be selected by the user to examine differences, for example, in cell lines with and without mutations in *KRAS*, or lung-versus liver-derived sample panels. The two per-gene likelihoods are then compared in a likelihood ratio test, with significance assessed by comparison with an empirical null distribution based on simulated data (see Methods). In this way, ACE performs a rigorous test of the alternative hypothesis that gene *G* exhibits different degrees of essentiality in the test and control categories against a null hypothesis of equal essentiality.

### Simulation Study

To evaluate the ability of ACE to infer sgRNA-, sample-, and gene-specific parameters, we devised a simulation method to generate artificial CRISPR-screen data sets. To ensure that our simulated data resemble data from real CRISPR screens as closely as possible, our simulator, called empiriCRISPR, mimics the variation in read counts observed in a reference experiment and makes minimal assumptions about the process by which the data are generated (see Methods and Supplementary Fig. S3). Importantly, empiriCRISPR is fully independent of the probabilistic model and parameter inference used in ACE.

### Benchmarking of Absolute Essentiality Predictions

Using empiriCRISPR, we generated data sets of 5,250 genes in which the essentiality of each gene ranged from complete nonessentiality (*ϕ_G_* = 0) to complete essentiality (*ϕ_G_* = 1). To ensure a fair comparison with methods that rely on a median-based normalization method, we took care to include an adequate number of nonessential genes in each data set, simulating 3,150 genes with *ϕ_G_* = 0 (60% of all genes). The remaining 40% of genes consisted of 300 genes at each of several intermediate values of *ϕ_G_* and 600 genes at *ϕ_G_* = 0.99 (see Fig. 2a). For each choice of simulation parameters, three replicates were simulated with four sgRNAs targeting each gene and an average of 300 reads per sgRNA. As a template data set for empiriCRISPR, we used a collection of previously published genome-wide CRISPR screens in which both the master library and initial infected sgRNA abundances were sequenced along with the depleted samples (16). The master library was generated through random selection of sgRNA from the template screen, followed by simulation of read counts according to the variation observed in the template.

**Fig. 2.**
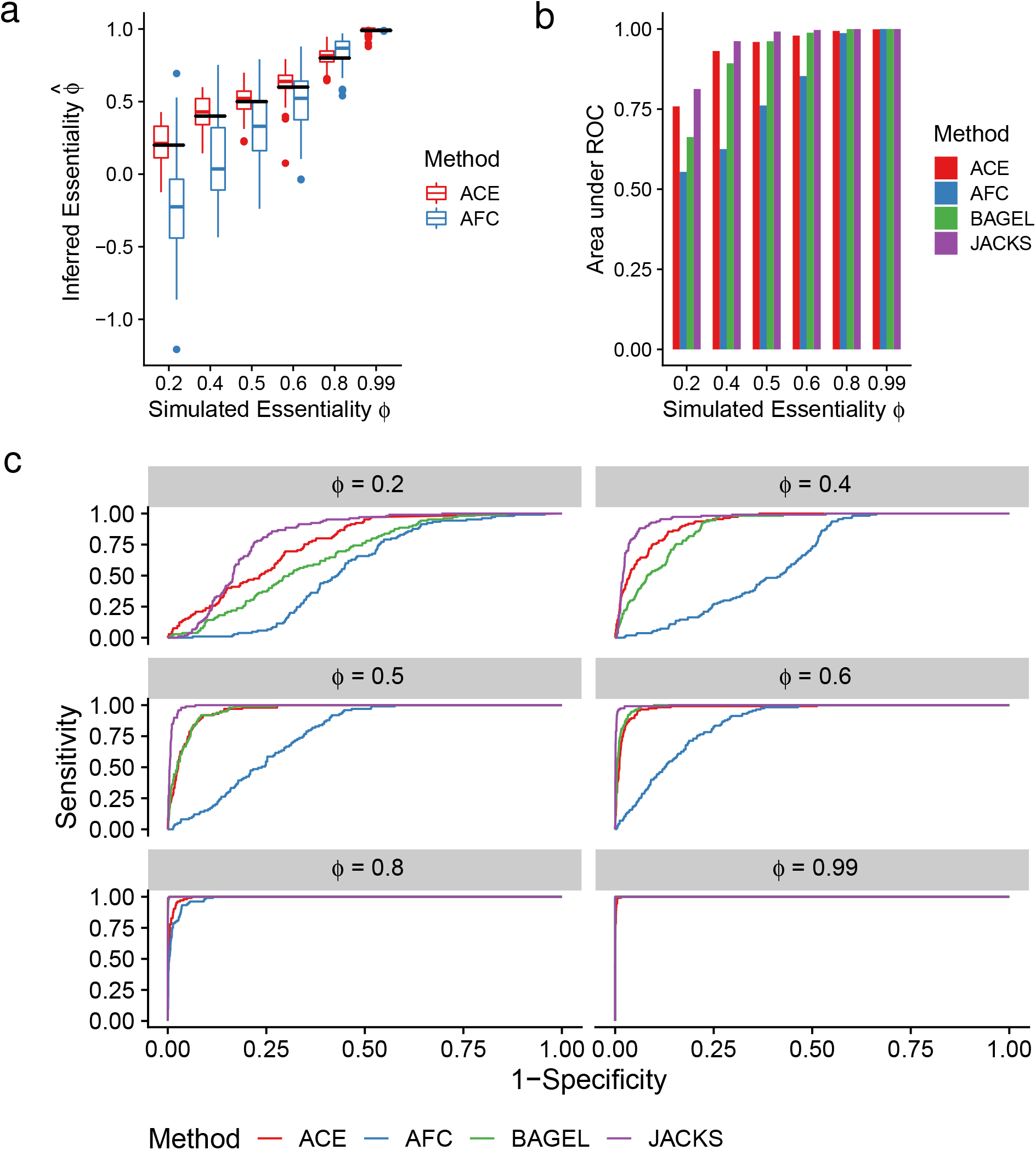
Detection of Absolute Essentiality in Simulated Data. (a) Inferred essentiality values 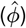 for 300 genes simulated in three replicates at each essentiality value shown (*ϕ*). Simulated data sets also included 3,150 nonessential genes (*ϕ_G_* = 0; not shown), 300 of which were provided to each method as negative controls. Black horizontal bars indicate true values 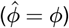. (b) Performance in binary classification (essential vs. nonessential) of simulations of 300 ‘essential’ genes at various values of *ϕ_G_* and 300 ‘nonessential’ genes. Performance reported as the Area under the Receiver Operating Characteristic (ROC) curve. Only initial and final read counts from the simulation were used for all methods. (c) Full ROC curves for each of the simulated essentiality values. ‘ACE’ — our probabilistic method (thresholded on log likelihood ratios), ‘AFC’ — method based on average fold changes in sgRNA abundance (thresholded on *z*-scores; see Methods). ‘BAGEL’ — Bayesian Analysis of Gene Essentiality (2) (thresholded on reported Bayes Factors); ‘JACKS’ — Joint analysis of CRISPR/Cas9 knockout Screens (8) (thresholded on reported *p*-values).

We first compared ACE’s estimates of essentiality with an estimator based on the average fold change (AFC) in read counts (see Methods), which has been used in several previous studies (17, 29) (see Fig. 2a). The ACE estimates are approximately unbiased and exhibit relatively low variance (see also Supplementary Fig. S4). By contrast, the AFC-based estimates display a downward bias, particularly at low simulated values of *ϕ_G_*, and show substantially greater variance. Notably, the simple AFC-based method often yields negative estimates of *ϕ_G_* even when the true *ϕ_G_* is positive.

Many methods do not provide a numerical estimate of the essentiality of a gene, but instead focus on a binary classification of genes into ‘essential’ and ‘nonessential’ categories. To compare ACE with these methods, we simulated data sets containing 300 genes at each of several values of *ϕ_G_*, together with 3,150 nonessential genes. For each of these experiments, we then compared the performance of ACE in binary classification with BAGEL (1), JACKS (8), and a classifier based on simply thresholding the AFC statistic. We summarized the performance of all methods using Receiver Operating Characteristic (ROC) curves and the Area Under the ROC Curve (AUC) metric (Fig. 2b and (c)). We found that all methods performed similarly well at high values of *ϕ_G_*, with the AFC-based method performing slightly worse, but in the harder scenarios where *ϕ_G_* takes moderate to low values (*ϕ_G_ <* 0.5), JACKS and ACE performed significantly better than the others. However, JACKS slightly outperformed ACE on this task, apparently because it makes particularly efficient use of the negative controls in defining a null distribution.

### Benchmarking of Differential Essentiality Predictions

To test the detection of differential essentiality, we simulated CRISPR-screen data in two samples (‘test’ and ‘control’) that included 300 genes at each of several values of 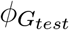 that were essential only in the ‘test’ sample (see Methods). To represent uniformly essential genes, both samples also included 300 genes at strong (*ϕ_G_* = 0.99) and 300 genes at moderate (*ϕ_G_* = 0.5) essentiality, as well as a large collection (3,150) of nonessential genes (*ϕ_G_* = 0). Similar to the previous tests, we compared the performance of ACE with BAGEL, JACKS, and a simple AFC-based method in the detection of the differentially essential genes. Because the other methods do not support direct tests for differential essentiality, we devised tests based on the results of two separate tests for essentiality. After some experimentation, we settled on using the ratio of *p*-values for the test and control panels for JACKS, and the absolute difference in the Bayes Factors for the test and control panels for BAGEL (see Methods).

We found that ACE’s likelihood ratio test for differential essentiality did result in substantially better power than the competing methods (Fig. 3). All methods had reasonably good power when 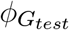 was large (*≥* 0.6) with JACKS performing somewhat more poorly than the others; but at values of 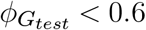, ACE showed a clear advantage. Even when 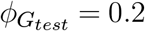, ACE was able to detect 67% of differentially essential genes (Fig. 3b).

**Fig. 3.**
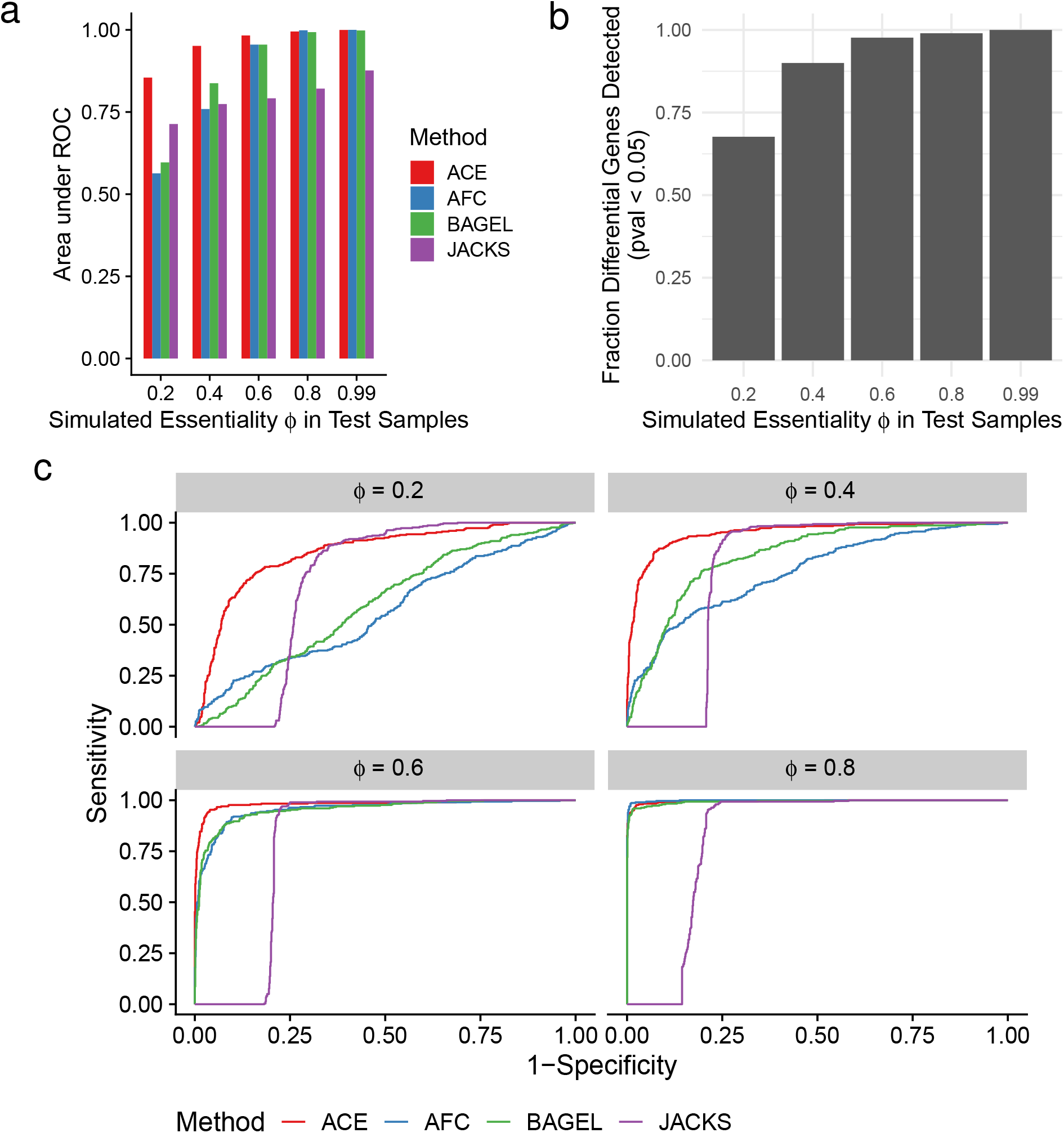
Detection of Differential Essentiality in Simulated Data. (a) Performance in binary classification of differential essentiality for 300 genes that were ‘essential’ (at various levels of simulated essentiality *ϕ*) in ‘test’ samples and ‘nonessential’ (*ϕ* = 0) in ‘control’ samples. Both samples also included 300 genes at strong (*ϕ_G_* = 0.99) and 300 genes at moderate (*ϕ_G_* = 0.5) essentiality, as well as a large collection (3,150) of nonessential genes (*ϕ_G_* = 0). 300 genes from each of these uniformly essential sets were used as negative controls. (b) Fractions of differentially essential genes correctly detected by ACE (at a Bonferroni-corrected empirical *p* < 0.05) at various values of simulated essentiality *ϕ*. (c) Full ROC curves for each of the simulated differential essentiality levels. In all cases, three replicates of the control and test samples were simulated for each value of *ϕ*. Methods are as described in Fig. 2.

## Analysis of Real Data

### Identification of Essential Genes

We next sought to evaluate the performance of ACE in detecting previously identified essential genes from real data. We focused on 688 genes that have been classified as essential in at least two previous studies (1–3), based on both RNAi and CRISPR screens (19). For negative controls, we used 634 genes that are not widely expressed according to public RNA-seq data (see Methods). We used ACE to evaluate the essentiality of both gene sets using the CRISPR-screen data from Achilles DepMap. Notably, this data set includes sequence data representing only the sgRNA master library and the post-depletion sgRNA abundances. To ensure a similar genetic background between samples, we focused on 92 unique cell lines that all originate from lung adenocarcinoma, a subtype of non-small cell lung cancer (NSCLC).

Considering the many differences in methods and data sets, we found that ACE’s characterization of these genes was fairly concordant with previous results. The genes previously identified as essential (positive controls) yielded both clearly elevated estimates of essentiality 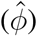 and clearly elevated log likelihood ratios (LLRs) in a test for non-zero essentiality relative to the negative controls (Fig. 4a). At a LLR threshold defined such that only 5% of the negative controls exceeded it (i.e., with empirical *p* = 0.05), 90.5% of the positive controls were predicted to be essential. In an ROC analysis based on the same data, ACE was able to distinguish the positive and negative controls with high accuracy (AUC = 0.97; see Supplementary Fig. S5).

**Fig. 4.**
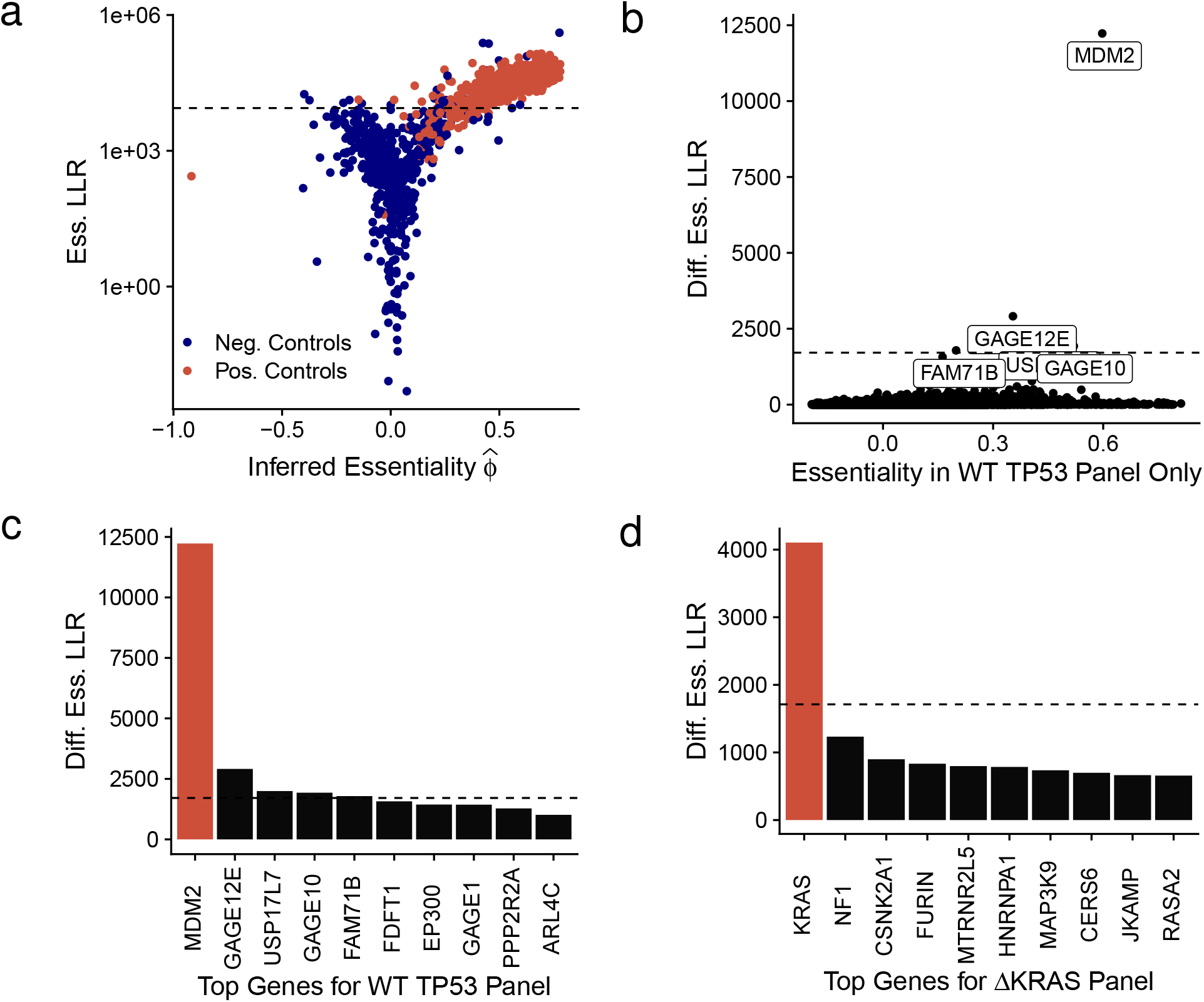
Identification of known genotype-specific essentiality. (a) Estimates of essentiality per gene (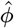; *x*-axis) and essentiality log likelihood ratios (Ess. LLR; *y*-axis) from ACE based on genome-wide CRISPR-screen data from the Achilles DepMap Project. Results are shown for 688 ‘essential’ genes (Pos. Controls; *red*) identified in references (1–3) and 634 nonessential genes (Neg. Controls; *blue*; see Methods for details). The LLR metrics are relative to a null hypothesis of 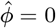 (b) Estimated essentiality per gene in wild-type *TP53* (*ϕ*; *x*-axis) vs. log likelihood ratio (LLR) for differential essentiality in wild-type vs. mutant *TP53* based on DepMap data for NSCLC Adenocarcinoma (9). Several high-scoring genes are highlighted with labels, including *MDM2* (see text). (c) Top 10 genes identified as differentially essential in NSCLC samples having wildtype *TP53*, ordered by differential essentiality log likelihood ratio (Diff. Ess. LLR). (d) Top 10 genes identified as differentially essential in NSCLC samples having mutant *KRAS*. In panels (b)–(d), the horizontal dashed lines indicate Bonferroni-corrected empirical *p*-values of 0.05. Genes highlighted in red in panels (c) and (d) are discussed in the text.

### Detection of Known Cases of Genotype-Specific Essentiality

One possible application of ACE’s test for differential essentiality is to identify significant differences in essentiality between distinct genetic backgrounds. For example, cases of synthetic lethality may appear as genes that are essential in test samples in which another gene is mutated and nonessential in control samples in which that gene is intact. Conversely, other genes may be found to be essential in the presence of the wildtype but not the mutant genotype. To explore this possibility, we contrasted cell lines annotated by the Cancer Cell Line Encyclopedia (CCLE) with variants of two very well-known cancer-related genes, the tumor suppressor *TP53* and the proto-oncogene *KRAS* (see Methods) (30, 31).

We restricted ourselves to NSCLC samples in all cases, to avoid spurious signals unrelated to the mutations of interest. Moreover, we structured our test specifically to identify genes that were significantly more (rather than less) essential in the test samples than in the control samples. As in the previous analysis, we used data from Achilles DepMap (9, 24).

In the case of *TP53*, we first defined the test samples as ones with ‘wildtype’ copies of *TP53*, with no annotated mutations, and the control samples as ones in which *TP53* has ‘damaging’ mutations (denoted here as Δ*TP53*; see Methods). This test for a wildtype-dependent increase in essentiality yielded a clear outlier: the gene *MDM2* (Fig. 4b and (c)). It is well known that cell lines with wildtype *TP53* require an intact copy of *MDM2* to negatively regulate the TP53 protein and allow for cell proliferation (32, 33). This result therefore provides support for the biological validity of our test for differential essentiality.

In the case of *KRAS*, which becomes oncogenic in the presence of gain-of-function mutations and is known to drive aggressive tumorigenesis (34), we defined the samples in the opposite way: with test samples containing a missense mutation in *KRAS* (Δ*KRAS*), the majority of which are activating gain-of-function mutations (35), and control samples with no annotated SNP (WT*KRAS*). Samples containing wildtype *TP53* were additionally excluded to remove the signal from *MDM2*. As predicted, the *KRAS* gene itself emerged as having, by far, the strongest signal for a mutation-dependent increase in essentiality (Fig. 4d), further validating our method.

### Test for Non-Oncogene Addictions in TP53 Mutants

Finally, we swapped the ‘test’ and ‘control’ samples for *TP53* — using Δ*TP53* samples for the ‘test’ and wildtype samples for the ‘control’ — with the goal of discovering novel non-oncogene addictions. As discussed above, such addictions may suggest new drug targets in cases of undruggable loss-of-function mutations. This test of increased essentiality in the presence of damaging *TP53* mutations identified several candidate genes (Fig. 5a and Supplementary Fig. S6).

**Fig. 5.**
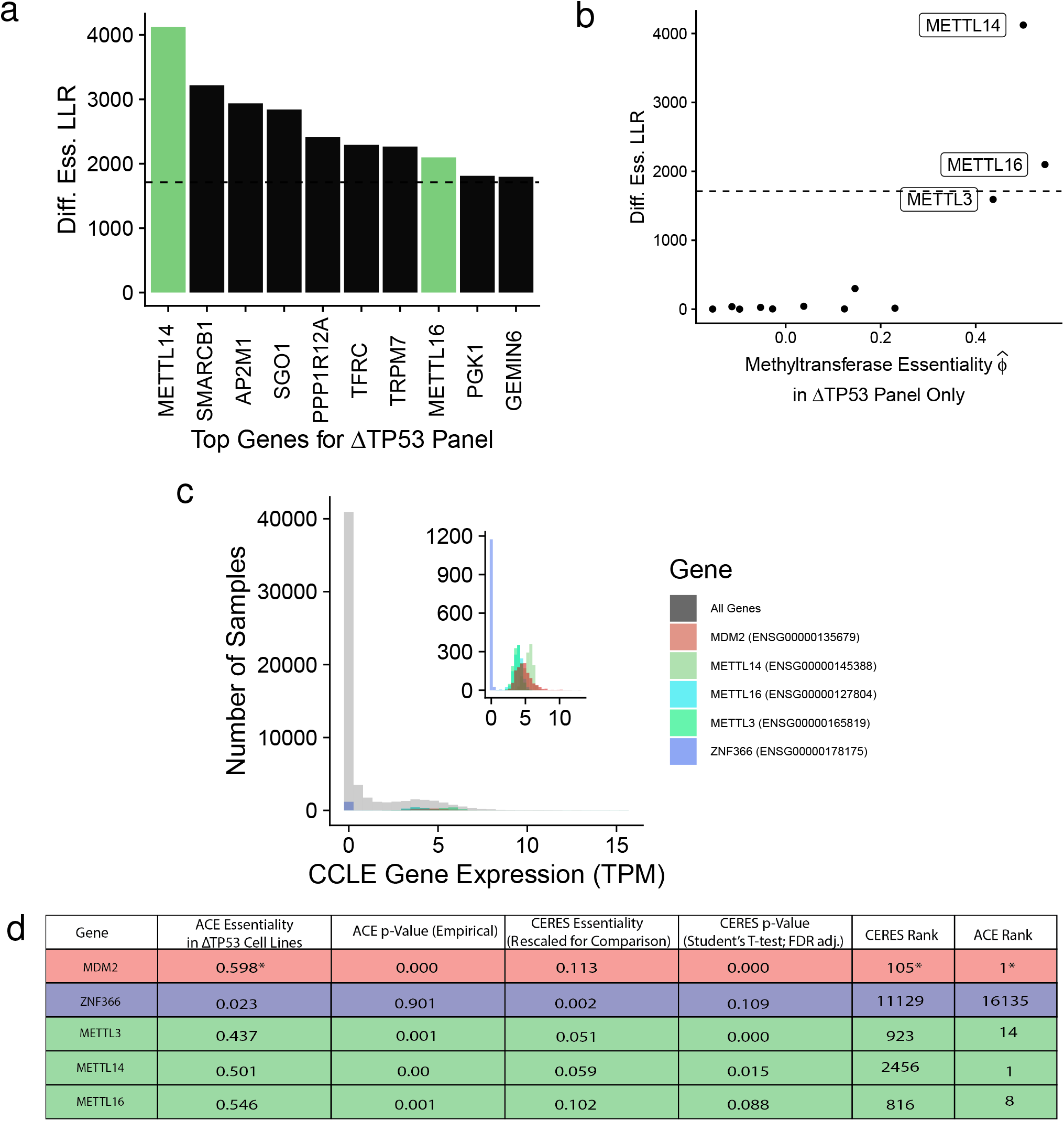
RNA Methyltransferase Genes Associated with***TP53***. (a) Top 10 genes (by differential essentiality log-likelihood ratio) showing increased essentiality in NSCLC cell lines that harbor ‘damaging’ mutations in *TP53* (Δ*TP53*) according to the CCLE, suggesting pos-sible non-oncogene addictions. The RNA m^6^-A methyltransferases *METTL14* and *METTL16* (*green*) are of particular interest (see text). (b) Estimated essentiality of RNA methyltransferase genes in Δ*TP53* samples (*x*-axis) vs. log likelihood ratio in Δ*TP53* vs. wild-type *TP53* samples (*y*-axis). (c) Gene expression data from the CCLE, processed by the Achilles DepMap, for the Δ*TP53* cell line panel highlighting the differentially essential methyltransferase genes from panel (b). *MDM2* is shown as a positive control and nonessential *ZNF366* is shown as a negative control. The distribution of all genes is shown in gray. (d) Comparison of differential essentiality candidates and controls based on ACE and the CERES method from Project Achilles (see Methods for details). An asterisk indicates a result applies for values associated with *MDM2* only in the presence of wildtype *TP53*; other genes were tested for essentiality specific to cell lines with a damaging mutation in *TP53* (ΔTP53). In panels (a) and (b), the horizontal dashed lines indicate Bonferroni-corrected empirical *p*-values of 0.05.

Notably, the top candidates included two genes encoding RNA m^6^-A methyltransferases, *METTL14*, and *METTL16*. A third gene, *METTL3*, encoding the heterodimeric partner of *METTL14*, also scored highly for differential essentiality (Fig. 5b). These three genes were assigned moderately high scores for differential essentiality according to the CERES method from the Achilles Project, but CERES did not rank any of them among its top 100 differentially essential genes (Fig. 5d). This difference in ranking between ACE and CERES possibly reflects ACE’s improved power for detecting differential essentiality. All three methyl-transferases are expressed at a moderate to high level across the *TP53*-mutant cell line panel, according to CCLE expression data (Fig. 5c and Supplementary Fig. S7) (24, 30, 31).

Several studies have noted CRISPR sgRNA cutting in multiple regions can reduce cell proliferation, regardless of the essentiality of the sites targeted. To ensure that the signatures of essentiality in *METTL3*, *METTL14*, and *METTL16* are not merely reflective of changes in copy number, we compared gene duplications between mutant and wildtype *TP53* sample panels (Supplementary Fig. S8). Copy number variation was similar between the two panels, suggesting that differences in gene essentiality are driven by biological activity of the genes themselves. In addition, flanking genomic regions of *METTL14* and *METTL16* do not contain known oncogenic drivers, eliminating the possibility that CRISPR-induced lesions are simultaneously influencing a nearby essential gene (Supplementary Fig. S9). Overall, our findings suggest that *METTL3*, *METTL14*, and *METTL16* are promising candidates for non-oncogene addiction in the absence of functional *TP53*.

## Discussion

In this paper, we have introduced a new probabilistic modeling and inference method, called ACE, for detecting essential genes and estimating levels of essentiality from CRISPR negative selection screens. As we have shown, ACE uses a hierarchical Bayesian model to account for the uncertainty associated with each major stage of a CRISPR screen and enable maximum-likelihood estimation of gene-level essentiality. It also supports likelihood ratio tests for both absolute and differential essentiality.

While the number of known essential genes is growing, simulations remain critical in providing an objective gold standard to evaluate the performance of computational methods. To test ACE and compare it with other available methods, we developed a new simulator, called empiriCRISPR. Our strategy with empiriCRISPR was not to generate simulated data according to a process-based model — which would tend not to account for all of the sources of noise in real data, and would be difficult to define without heavily overlapping the assumptions of ACE itself — but instead to mimic the empirical properties of real data sets as closely as possible. In this way, we were able to generate ‘authentic-looking’ data sets of arbitrary size with known sets of essential genes at known levels of essentiality. We showed that ACE performs quite well on these data sets despite their realistic ‘noise’ characteristics.

To our knowledge, ACE is the first method to support a direct likelihood-based test for differential essentiality between designated ‘test’ and ‘control’ sample panels. This test can be useful in a variety of applications, including tests for differences in essentiality in the presence of mutant and wild-type versions of known cancer genes. In comparison to indirect tests for differential essentiality — derived from separate analyses of ‘test’ and ‘control’ panels — ACE appears to offer improved power according to our simulation study (see Fig. 3). Notably, it is straightforward to define one-sided variants of this test to detect *increased* or *decreased* essentiality in the ‘test’ relative to ‘control’ panels. The framework also extends naturally to tests of more complex hypotheses (e.g., test essentiality *>* 0 and control essentiality *≤* 0). According to standard statistical theory, these will all be well-powered tests, provided they are applied to sufficiently abundant data for the asymptotic properties of likelihood ratio tests to become apparent.

As illustrated by the major revisions of ‘core essential gene’ sets over the past 10 years, the definition of ‘essential genes’ is inevitably somewhat subjective and arbitrary. In reality, essentiality occurs on a spectrum and depends on the particular genetic background, cell-type, and conditions tested. As a result, candidate ‘essential’ genes require carefully designed experiments for validation, and researchers should be cautious about relying upon published sets of genes labeled ‘essential’. Estimates of *ϕ_G_* in ACE are also limited, in that they reflect an ‘average’ essentiality across all samples analyzed, so users must take care to choose a panel of samples that ensures a level of diversity appropriate for the biological question of interest. For example, a diverse collection of tissue samples or cell lines may be most suitable for the identification of universally essential genes, whereas a set of patient-derived tissue samples may be best for personalized tumor therapeutics. Notably, because it is a relative metric, differential essentiality may in some cases generalize better than absolute essentiality. We anticipate that findings of differential essentiality may be of interest to researchers exploring genetic vulnerabilities even in other tissues and cancer subtypes.

Among the genotype-dependent differentially essential genes identified by ACE, we found the RNA methyltransferase genes *METTL3*, *METTL14*, and *METTL16* to be particularly interesting. RNA m6-A modifications have recently been found to be critical in the maintenance of leukemic cell growth (36). These genes have been previously implicated in other cancer types, and RNA m6-A methyltransferases in general have been a topic of recent interest in the new field of cancer epitranscriptomics, with several phase I trials targeting METTL3 due to begin in 2021 (21). METTL3, in particular, was recently identified as interacting with *TP53* mRNA and inducing drug resistance in a biochemical assay in SW48 colon cancer cells (37). Another study found *METTL3* expression to be essential in NSCLC adenocarcinoma, although only two cell types were examined (38). *METTL3* mRNA knockdown with siRNA has also been shown, by M6A-seq, to result in an enrichment of methylation changes in genes in the p53-signaling pathway (39). Finally, patients with genetic alterations of m6A regulatory genes have been found to have an increase in *TP53* mutations, albeit only with sample sizes of *n* = 5 (40). Thus, there is abundant indirect evidence suggesting that these genes deserve further investigation as possible candidates for addiction in the absence of functional *TP53*. More generally, we anticipate that improved tests for differential essentiality, such as the ones introduced here, combined with rapidly growing CRISPR-screen data sets, will enable the detection of many new cases of genotype-dependent differences in essentiality, including both synthetic lethality and non-oncogene addiction.

## Methods

### Initial Infection

The initial abundance *n_sg_* of each sgRNA *g* within sample *s* is assumed to be given by a Poisson distribution with mean *µ*_1_ = *c_s_m_g_*, where *c_s_* is the total number of cells infected in the screen and *m_g_* is the relative frequency of *g* in the master library:

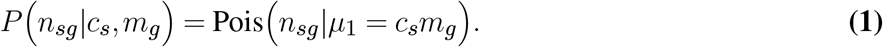

The value *m_g_* for each sgRNA *g* is estimated as the fraction of sequence reads corresponding to *g* in the sequence data for the master library. If multiple replicates are available, an average is taken. The parameter *c_s_* is either provided by the user or a default value of 1000 is assumed.

### Initial Read Counts

The sgRNA sequencing counts *x_sg_* from the initial infected population *n_sg_* are assumed to follow a Poisson distribution with mean *μ*_2_ = *γ_s_n_sg_*, where *γ_s_* is a scaling parameter that captures the relationship between sequencing depth and the number of cells infected:

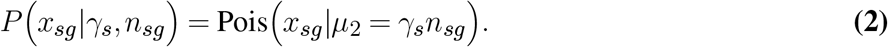

### Final Read Counts

Following several rounds of cell division, the final abundance of each sgRNA *g* is assumed to be *d_sg_* = *n_sg_*(1 *− ϵ_g_ϕ_G_*), where *ϕ_G_* is the essentiality of the gene *G* that is targeted by *g*, and *ϵ_g_* is the efficiency of *g*. This process is assumed to be deterministic; we assume that the number of infected cells is sufficiently large and sufficiently many cell divisions have occurred that stochastic effects can be ignored, and uncertainty in the value of *d_sg_* can absorbed by the distributions for *n_sg_* and *y_sg_*. The final read counts *y_sg_* are then assumed to follow a Poisson distribution with mean 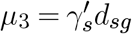, where 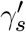 is the final read count scaling factor:

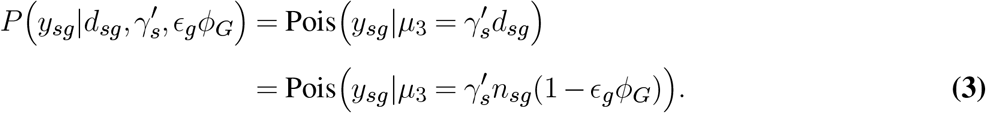

The scaling factor 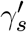 not only captures the relationship between sequencing depth and the number of cells that remain after cell growth, but also can accommodate sample-specific differences in the growth rate of all cells (independent of sgRNA).

### Guide-Specific Effects

The sgRNA-specific efficiency *ϵ_g_* is derived from a general feature vector by lo-gistic regression:

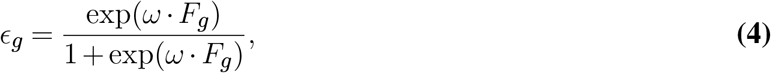

where *F_g_* is a feature vector and *ω* is a corresponding vector of real-valued coefficients. In this paper, we used a ten-dimensional vector *F_g_* whose elements are indicators (via “one-hot” encoding) for the decile of GC content of each sgRNA’s template guide. The corresponding 10-dimensional weight vector *ω* was treated as a free parameter and estimated by maximum likelihood. In this way, we effectively estimate a separate efficiency value for each GC decile, based on pooled information from all guides within that decile. As noted in the text, users may instead wish to use richer feature vectors such as those described in other recent studies (25, 41–45).

### Estimation of Sample Scaling Parameters

Following the ‘median-of-ratios’ approach used by DESeq2 (5), the sample scaling parameters *γ_s_* and 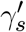 are not treated as free parameters in the maximization of the likelihood, but instead are set in a preprocessing step based on the raw read counts *x_sg_* and *y_sg_* for negative controls. Specifically, the scaling parameter is set equal to the median of the ratios of each sgRNA’s read count to a guide-specific reference value, which in this case is the expected number of infected cells, *c_s_m_g_*:

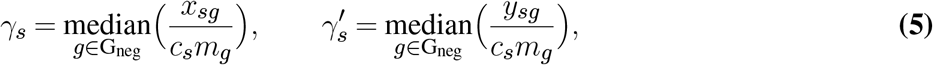

where *g ∈ G*_neg_ indicates all sgRNAs targeting genes in the designated set of negative controls. If no master library is available, the initial abundance of sgRNA 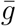 is approximated from the initial read counts as the average across samples:

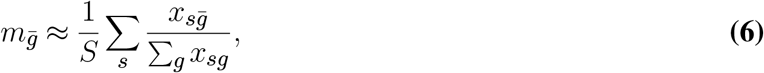

where *S* is the total number of samples. The user may choose to use the median ratio across all genes rather than only the negative controls, if desired.

### Full Model Likelihood

Substituting from equations 1–3, the full likelihood function can be expressed as:

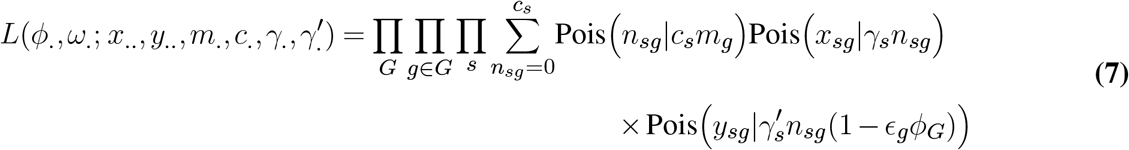

where *ϵ_g_* is implicitly a function of *ω* as defined by equation 4. Here and below we use “dot” notation to indicate sets of all relevant indices (e.g., *x_.._* = {*x_sg_* | for all samples *s* and sgRNAs *g*}).

The likelihood function must sum over the number of infected cells *n_sg_*, which cannot be directly observed. This summation step can be accelerated by an approximation that bundles summands together, for example, into groups of 10 cells. Weight vector *ω* is initialized to treat all sgRNA as 100% effective for the first optimization of *ϕ_G_*.

When *ω* is held fixed, the full likelihood decomposes by gene, such that each *ϕ_G_* can be estimated sepa-rately; and similarly, when the *ϕ_G_* values are held fixed, the likelihood decomposes by the efficiency features *f*, owing to our “one-hot” design for the feature vector. Thus, we estimate each *ϕ_G_* and component of *ω_f_* iteratively, to enable parallelization by gene and feature category. Specifically, we repeatedly carry out the following optimizations until convergence (for each gene *G* and feature *f*, respectively):

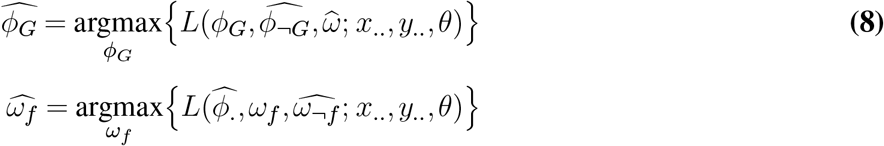

where 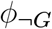 denotes the set 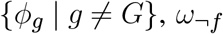 denotes the set {ω*_f′_* | *f′* ≠ *f*} and *θ* denotes the remaining parameter set.

Finally, to evaluate whether an estimated value of 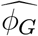 is significantly different from zero, we compare the component of the maximized likelihood function for that gene to the corresponding component when *ϕ_G_* is held fixed at zero. Specifically, we compute a test statistic *T* as,

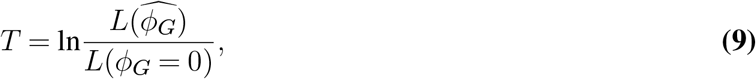

where *L*(*ϕ_G_*) represents the component of the likelihood corresponding to gene *G*. We compute an empirical *p*-value for *T* based on the equivalent test statistics for the designated negative control genes.

### Adaptation for Alternative Experimental Designs

The likelihood function can easily be adapted to settings in which data is unavailable for either the master library or the initial sgRNA abundances. In the case where the master library information is missing, we simply use a uniform prior for values of *n_sg_*, 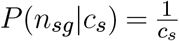 When the initial infected sgRNA abundances are unavailable, we remove the corresponding portion of the likelihood function, effectively treating *x_.._* as missing data:

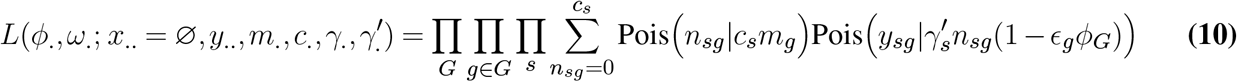

### Test for Differential Essentiality

To identify genes with significant differences in essentiality between a ‘test’ set of samples, *S_t_*, and a ‘control’ set, *S_c_*, we employ the following likelihood ratio test. Let *L*(*ϕ_G_*) represent the component of the likelihood function relevant to gene *G* that has been maximized in the standard way, with one version of the *ϕ_G_* parameter shared across both *S_t_* and *S_c_*. By contrast, let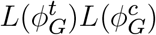 represent the corresponding component of the likelihood optimized such that separate versions of the essen-tiality parameter, 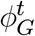 and 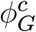, are used for samples *S_t_* and *S_c_*, respectively. The likelihood ratio test statistic *T*′ is then given by,

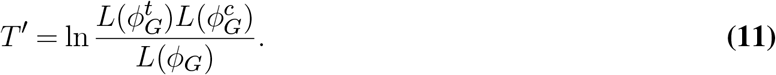

For this test, all samples are used to calculate the sample-specific parameters *γ_s_* and 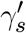, and the sgRNA-specific parameter *ϵ_g_*; only 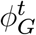 and 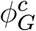 are re-estimated to compute *T′*. As with the test for absolute essentiality, an empirical *p*-value is computed for *T′* based on the distribution of values computed for the negative controls. In some analyses, we further require that 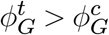 and otherwise set *T′* to 0.

### Essential and Nonessential Controls

The set of essential genes used for validation was defined as those genes that were included in at least two of three previously published sets of essential genes (1–3). The set of nonessential genes was obtained from 604 total RNA-seq datasets for 128 cell lines, in vitro differentiated cells, primary cells, and tissues accessed from the Encyclopedia of DNA Elements (ENCODE) data portal on August 29, 2019 (see Supplemental Data Table 1 for accession info of all cell lines used). For the nonessential set, we chose genes expressed at *≤* 1 TPM in all cell lines.

### Data from Cancer Dependency Map (DepMap)

For our analysis of real data from the Cancer Dependency Map (DepMap) Consortium, we focused on genome-wide CRISPR-screen data from Project Achilles, produced by the Broad Institute (9). This dataset uses the Avana library and includes 1374 publicly available screens at time of writing. Only the pre-infection master library and the post-depletion sgRNA abundances were sequenced, with one to four replicates per cell line available.

To retain a similar genetic background between sample panels, samples were compared from the same tissue type and lineage, specifically cell lines originating from lung Non-Small Cell Lung Cancer (NSCLC) Adenocarcinoma. The Achilles Project dataset includes 220 replicates of 92 unique samples of these cell lines. Following the Achilles DepMap convention (46), we assume a gene has lost function (indicated as

Δ) in all CCLE cell lines annotated as containing mutations that introduce single nucleotide polymorphisms (SNPs), deletions, or insertions in the start codon; deletions or insertions that introduce a frameshift mutation; mutations that introduce a premature stop codon or a *de novo*, frameshifted start codon; or mutations that disrupt a splice site (30, 31).

ACE results were compared with CERES published *p*-values available for each Achilles DepMap sample. As in ref. (9), the CERES differential essentiality test statistic was calculated using a one-sided Student’s *t*-test to compare the by-sample *p*-values of the test and control sample panels, and then was corrected by the rank-based false discovery rate.

### empiriCRISPR Simulation of CRISPR Knockout Screens

empiriCRISPR simulates each gene using a series of gamma distributions to represent each stage of a CRISPR knockout screen (infection, initial and final sequencing). CRISPR knockout screens from (16) were used as the empiriCRISPR template, and initial master library abundances were selected by sampling the template master sgRNA library. empiriCRISPR adjusts the gamma distribution parameters by minimizing the adjusted mean squared error of the summary statistics of simulated data with the median of 5 summary statistics of the template screens, shown in Fig. S3. The abundance of each sgRNA construct was assumed to be independent from all other sgRNA used in the experiment. Unless otherwise noted, simulations used 300 genes per essentiality value, 4 sgRNA per gene, and 3 samples.

### Package Software

ACE is implemented as an R package (R 3.5.0, (47)) using the *data.table* (48) and *R6* (49) packages. Optimization is performed using R’s *optim* package with the ‘Brent’ method. Both the ACE and empiriCRISPR github repositories can be found at https://github.com/CshlSiepelLab.

### Estimation of Essentiality from Mean Log-Fold Change

The ‘average fold change’ (‘AFC’) score for each gene *G* was calculated as follows:

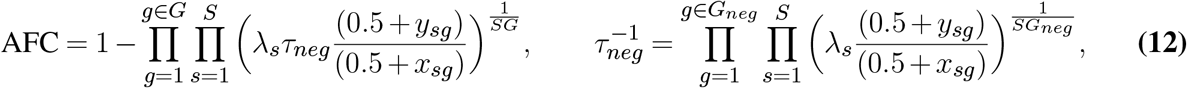

where a pseudocount of 0.5 was added to both initial (*x_sg_*) and final (*y_sg_*) read counts to prevent undefined values. Each sample was normalized by *λ_s_* to bring the total read count of each sample to a total of 10 million reads, and final fold changes were adjusted according to *τ_neg_*, the mean fold change in all negative controls *G_neg_*. Consistent with our ACE notation, an AFC value of zero has no impact on cell growth or proliferation.

### Estimation of Essentiality using BAGEL

We applied the BAGEL (Bayesian Analysis of Gene Essentiality) method (1) using software version 0.91 (last modified 09/2015). Simulated data was analyzed using 4% of genes as positive and negative controls, with simulated essentiality scores of 0.99 (99% depletion) and (5% depletion) respectively. Differential essentiality was calculated as the difference of Bayes Factors between separate analysis of test and control sample panels, as performed in (1). For the analysis of real data, we used all genes identified as essential or nonessential in our earlier analysis.

### Estimation of Essentiality using JACKS

We applied the JACKS (Joint Analysis of CRISPR/Cas9 Knock-out Screens) method (8) version 0.2 (downloaded 11/2019). Differential essentiality was evaluated by running JACKS separately for the test and control sample panels, using the built-in hierarchical prior option and the same negative control genes provided to other methods for both real and simulated data. The test statistic for differential essentiality was indicated by the absolute difference in *p*-value between test and control panels, as used in (8).

## Acknowledgments

We thank Professors Molly Hammell and Justin Kinney for advising on this project, Professor Yi-fei Huang for providing feedback on the mathematical design, and members of the Vakoc and Siepel labs for valuable advice and suggestions. Funding was provided by the Watson School of Biological Sciences William R. Miller Fellowship (to ERH), and the US National Institutes of Health grant R35-GM127070 (to AS). The content is solely the responsibility of the authors and does not necessarily represent the official views of the US National Institutes of Health.

## Supplementary Section 1: Supplemental Figures

**Supplementary Figure S1.**
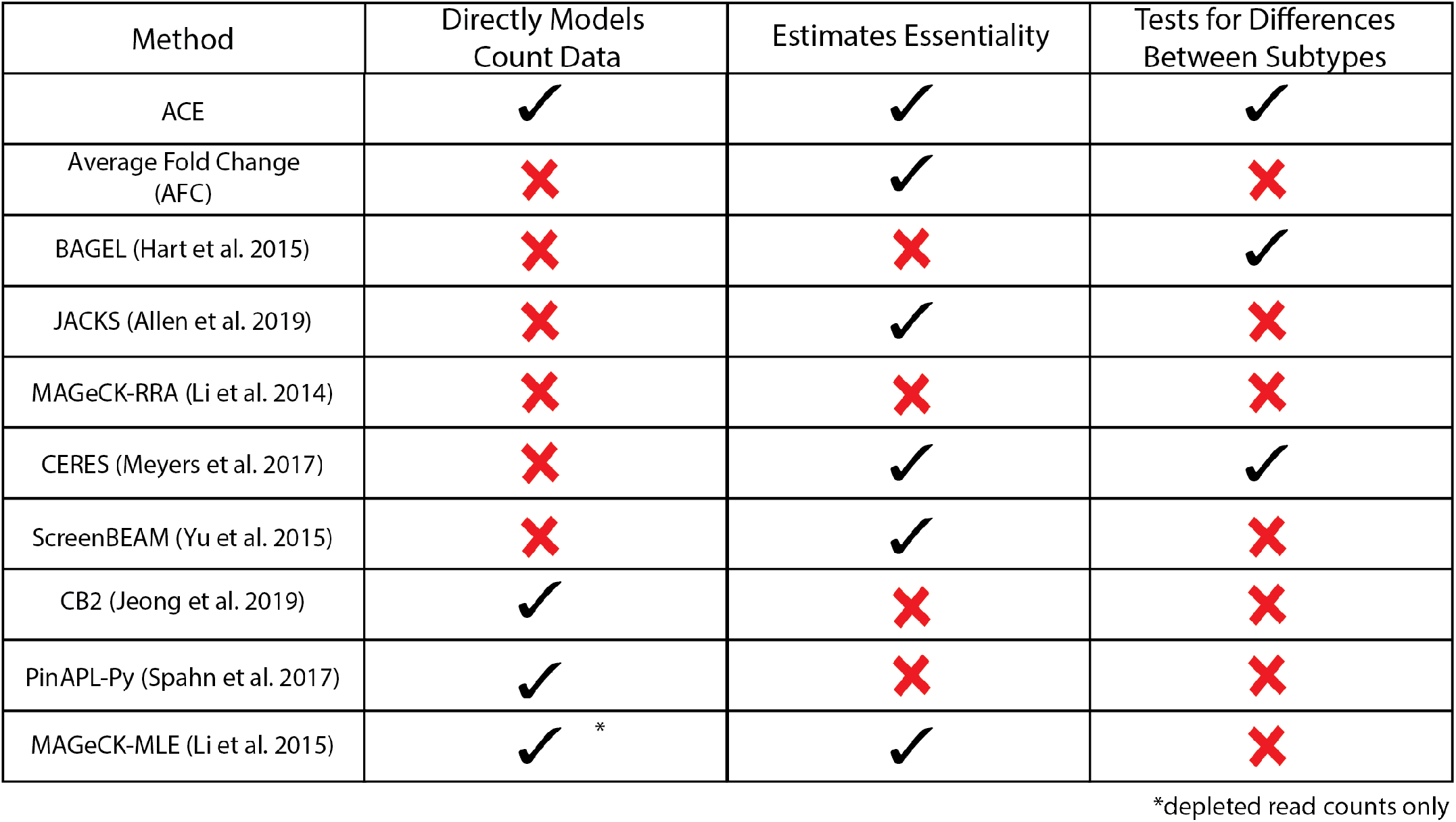
Methods for Identifying Differential Essentiality in CRISPR Screens. Many methods have been developed to analyze CRISPR screening data, but none are designed to test for differential essentiality between sample subtypes as a part of their inference framework. Some, such as BAGEL and CERES, test for significant differences in essentiality after parameter inference, but do not test whether the data is best described by a single essentiality parameter (‘Tests for Differences Between Subtypes’ column). Directly modeling count data, as opposed to relying upon summary statistics such as log-fold change, enables ACE to adapt to variations in experimental design, and leverage information from all available data (‘Directly Models Count Data’ column). Providing a quantification of essentiality, as opposed to fitting a binary essential/nonessential classification, enables researchers to more intuitively estimate phenotype (‘Estimates Essentiality’ column).

**Supplementary Figure S2.**
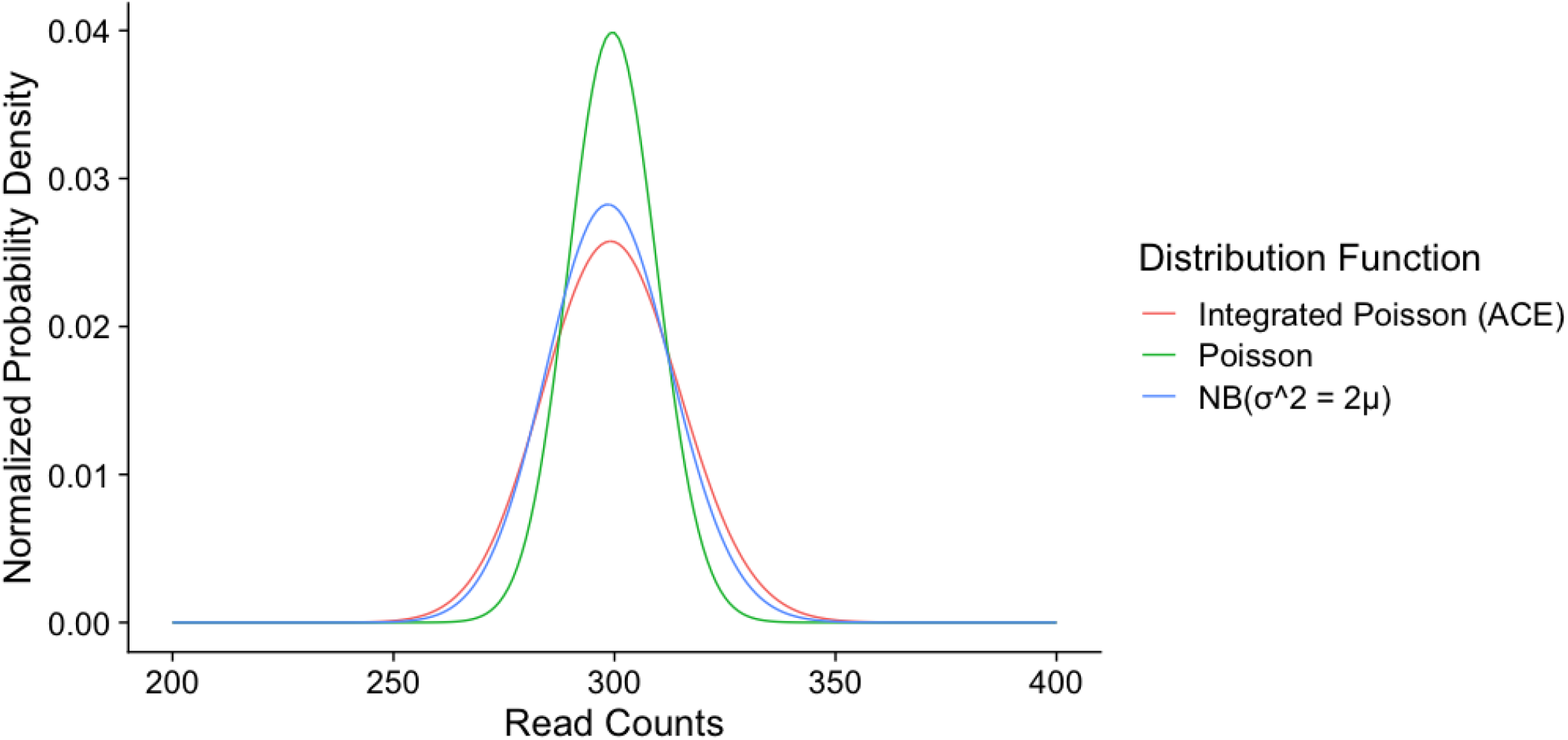
Illustration of Dispersion of ACE Distribution Function. Read count data is known to have a high dispersion similar to a Negative Binomial distribution (5). The values shown here reflect the probability density of different values of read counts in a CRISPR screen, as described by a Poisson distribution with point estimates of the mean (green), a Negative Binomial distribution with point estimates of the mean and a variance equal to twice the mean (blue). The ACE model integrates over all possible numbers of infected sgRNA, resulting in a probability density (red) similar to the Negative Binomial.

**Supplementary Figure S3.**
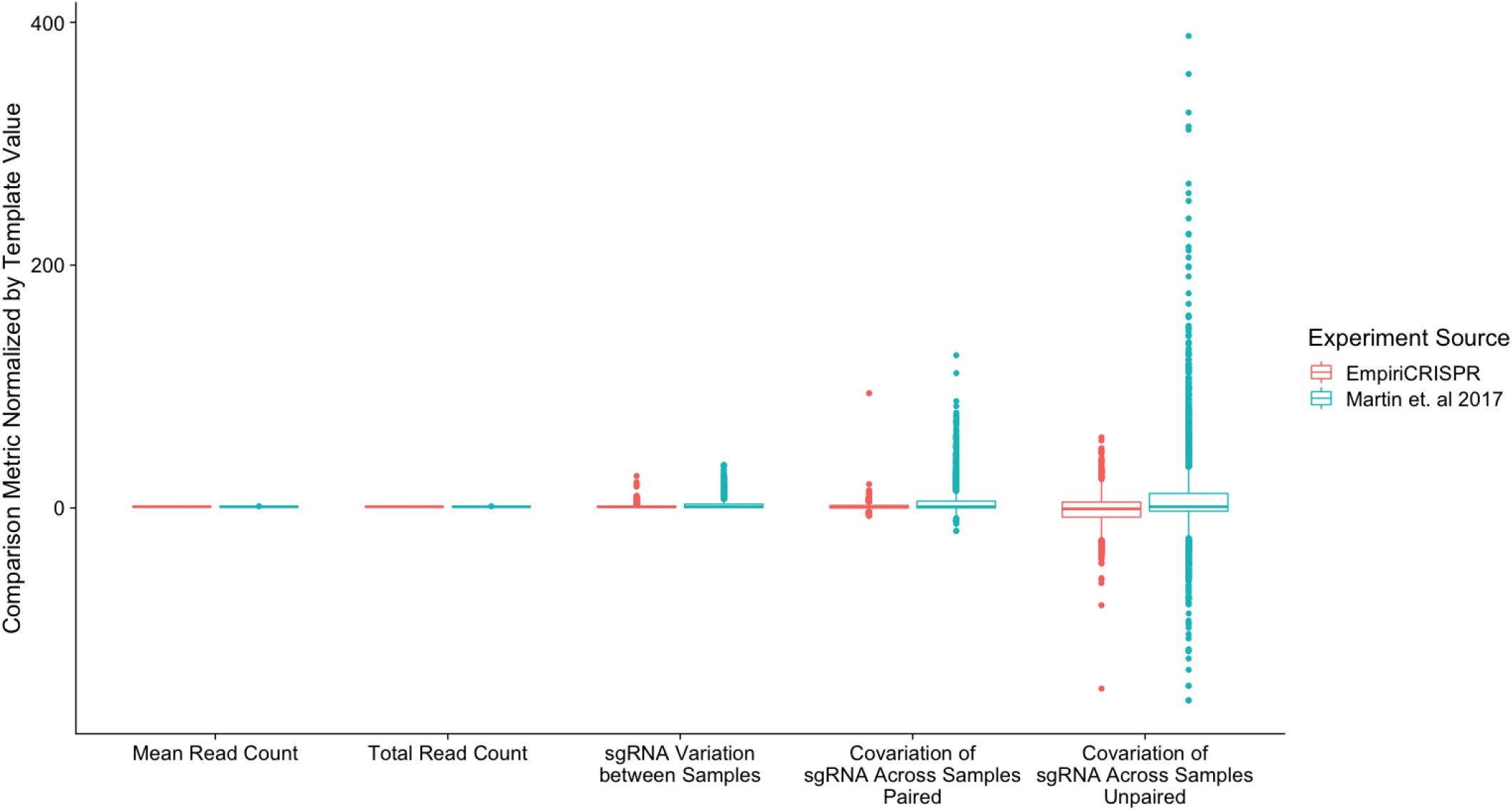
Simulated Data for Evaluation of ACE Performance. The genome-wide CRISPR screen performed by Martin et. al 2017 (16) was used as our EmpiriCRISPR template, as this work sequenced the sgRNA master library used as well as the initial and final samples. The sequenced master library provided the initial sgRNA abundances in the simulation, and all metrics shown were calculated in the initial and depleted counts. Shown are approximately 2300 guides targeting globally nonessential genes, as identified using ENCODE expression data (see Methods), and a randomly selected matching subset of nonessential sgRNA from simulated read counts. Mean and total read counts were calculated within each sample, while variation and covariation metrics were calculated for each guide across samples. ‘Paired’ indicates covariance was calculated between initial and depleted samples, with each sgRNA paired with the depleted sample produced by the same master library transfection; ‘Unpaired’ have been paired with samples from a different transfection replicate.

**Supplementary Figure S4.**
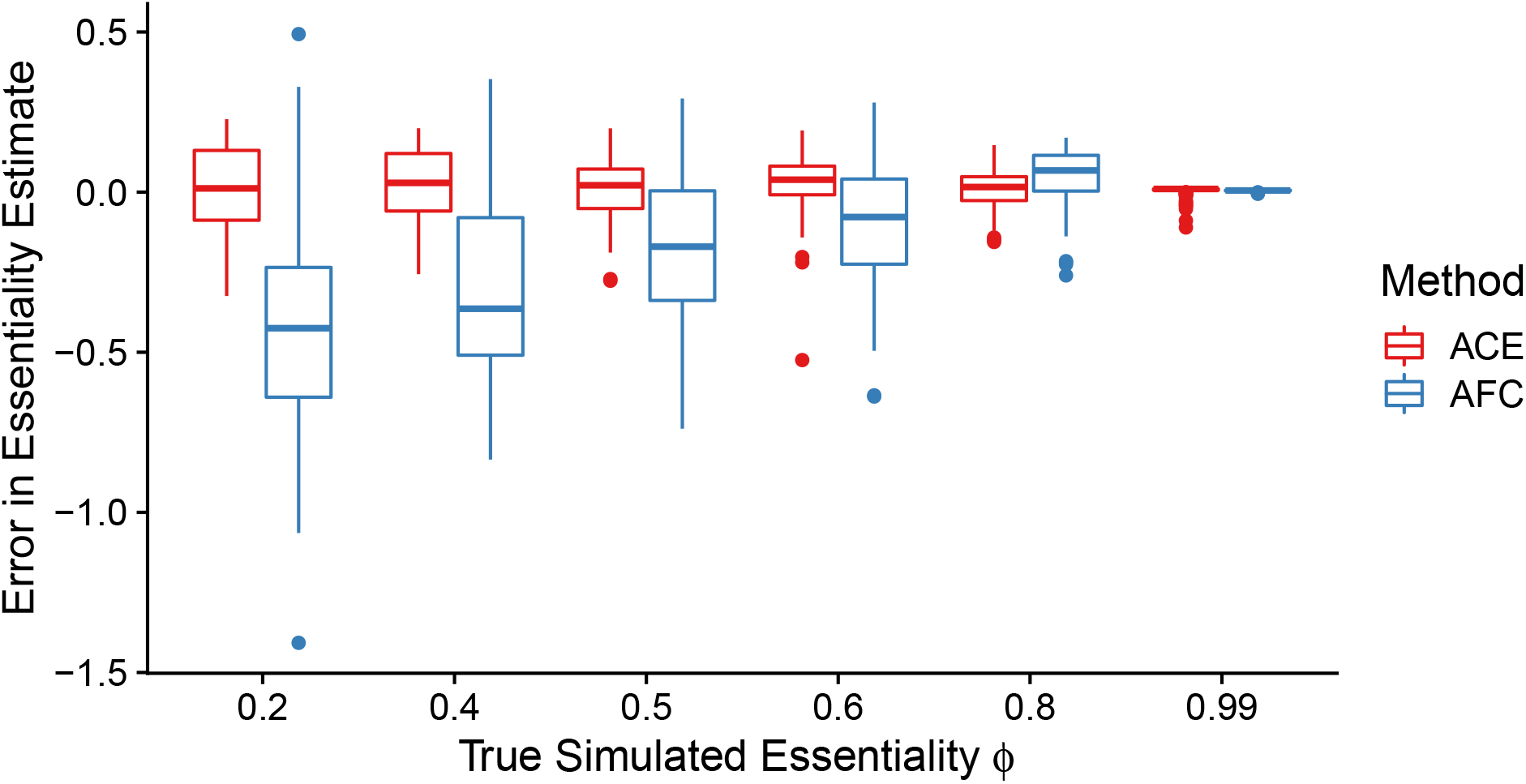
Error in Estimation of Gene Essentiality. Gene essentiality *ϕ_G_* was calculated using ACE and average fold change (AFC; see Methods) across three replicates of simulated count data. 300 genes were simulated at each essentiality level shown; results from an additional 3,150 nonessential genes are not shown.

**Supplementary Figure S5.**
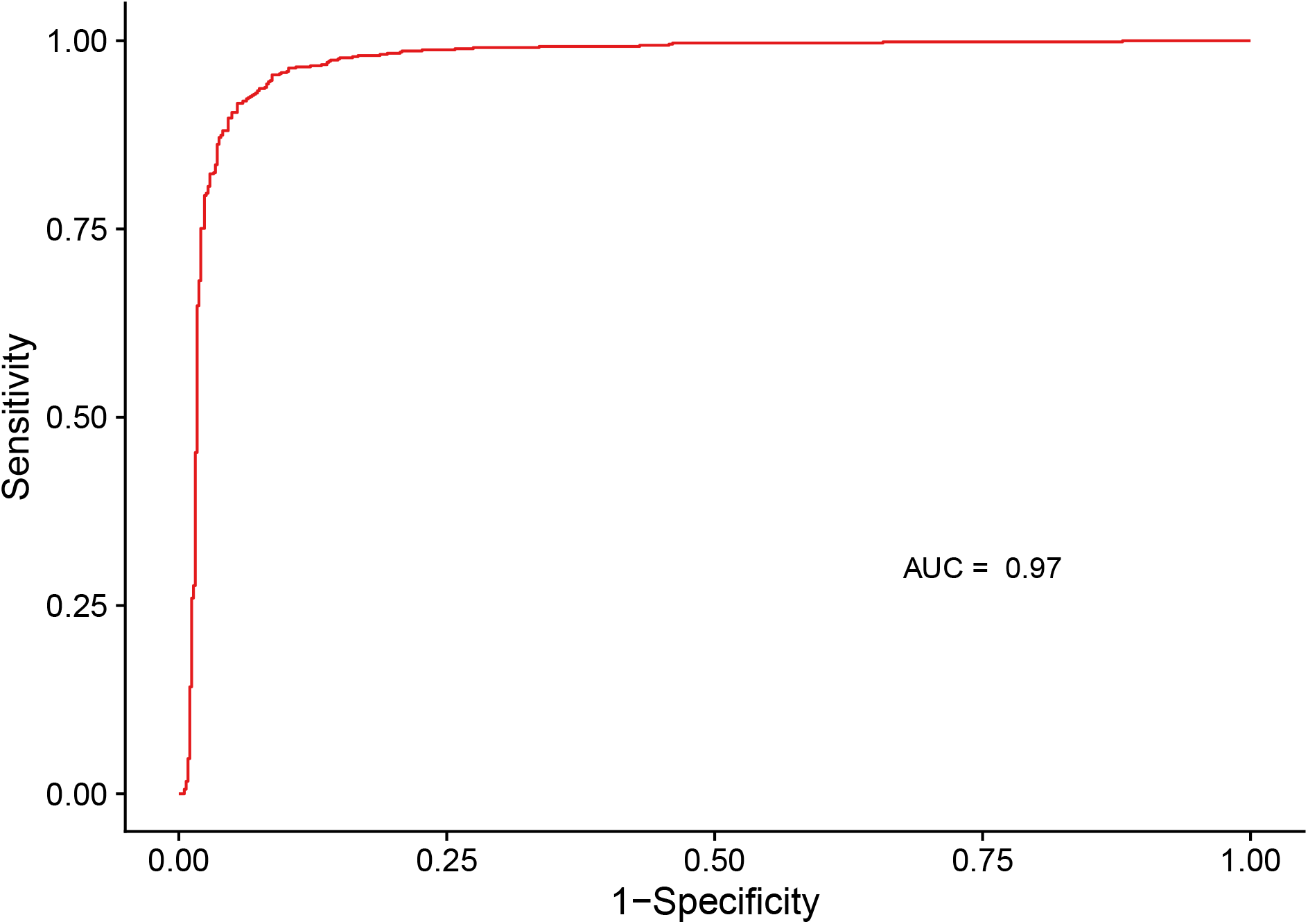
ACE Classification of Established Essential and Nonessential Genes in Achilles DepMap CRISPR Screens. All 220 CRISPR screens performed in 92 different NSCLC Adenocarcinoma cell lines in the Achilles DepMap project (9) were used by ACE to estimate the likelihood of essentiality of 688 essential genes.

**Supplementary Figure S6.**
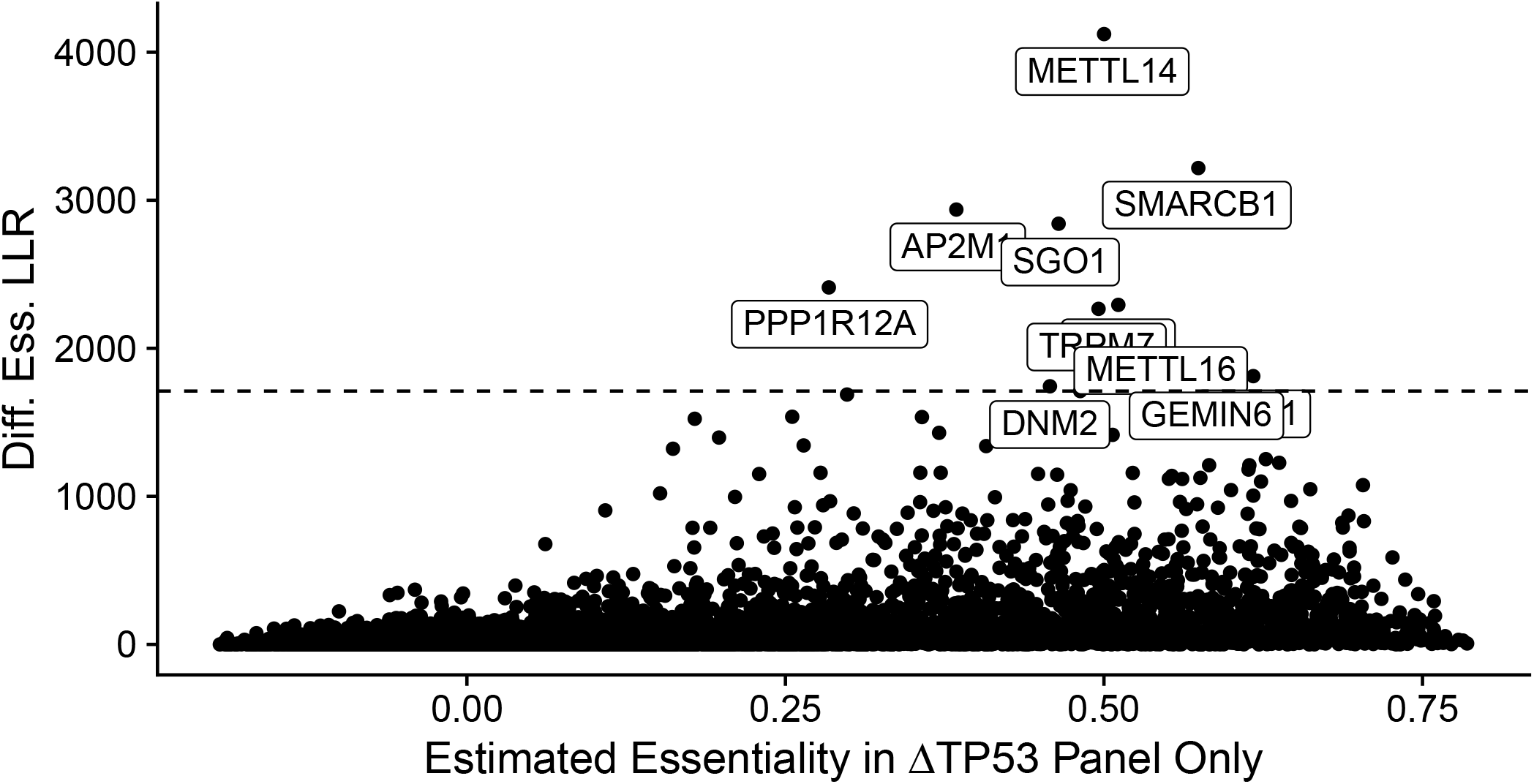
Genes Essential in NSCLC Adenocarcinoma cell lines with a ‘damaging’ mutation in *TP53*. All genes evaluated by genome-wide CRISPR screens in NSCLC cell lines by the DepMap Project Achilles are shown. On the x-axis is the ACE estimate of essentiality in cell lines with a nonsense or frameshift mutation indicated as ‘damaging’ in *TP53* (ΔTP53 Panel; see Methods). On the y-axis is the test statistic indicating whether this essentiality is significantly different from cell lines with wildtype *TP53*, calculated with a log likelihood ratio test comparing a shared versus a unique essentiality between the two sample panels. The horizontal line indicates the Bonferroni-correct empirical *p*-value <0.05.

**Supplementary Figure S7.**
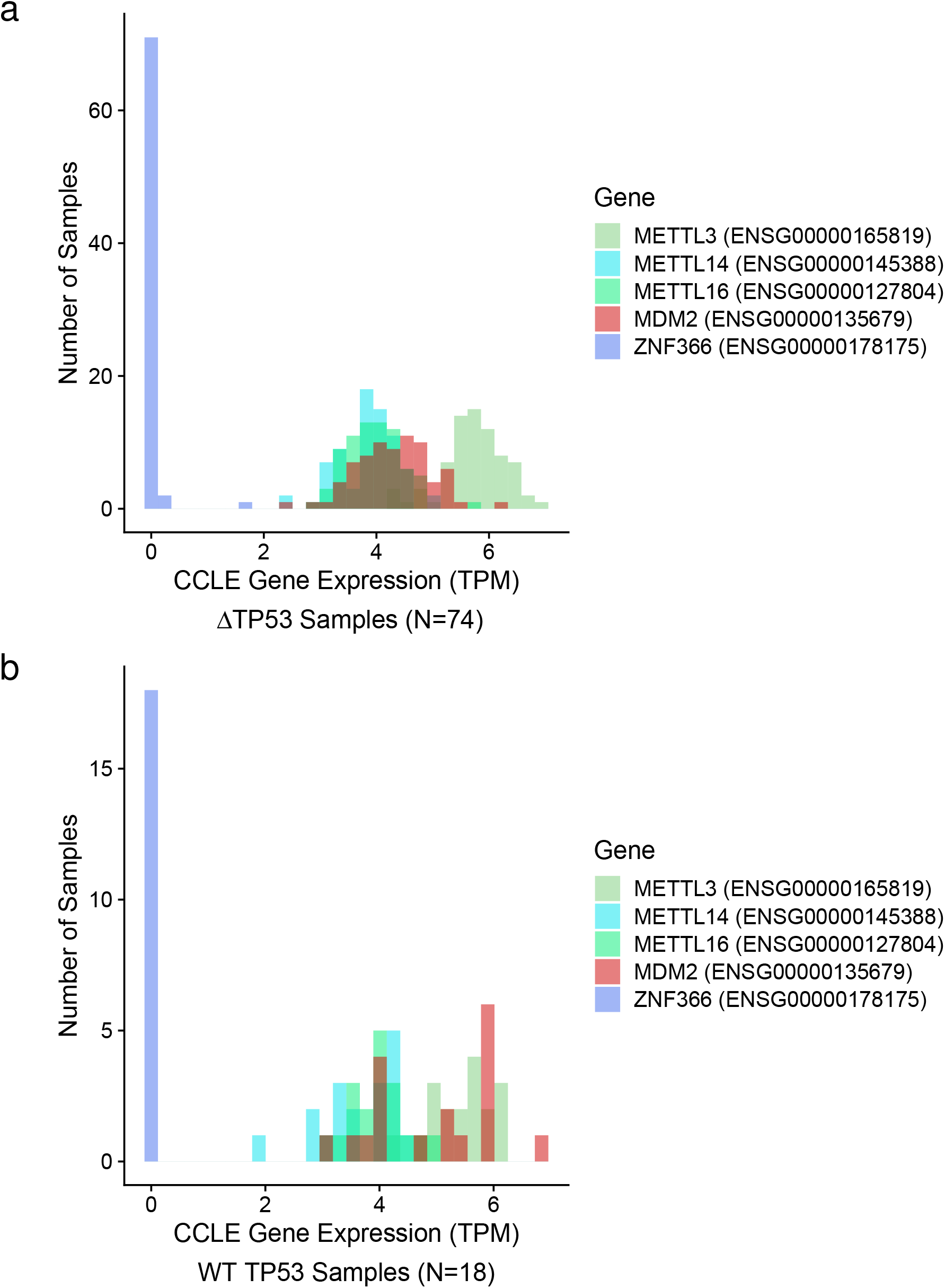
Expression Levels of Differential Essentiality Candidates. CCLE reported expression levels of differential essentiality candidates *METTL3*, *METTL14*, and *METTL16* in cell lines used by the Achilles DepMap project with annotated nonfunctional (a) and presumed functional (b) variants of *TP53* (9, 30, 31). *MDM2* is shown as a positive control for gene essentiality; *ZNF366* as a negative control.

**Supplementary Figure S8.**
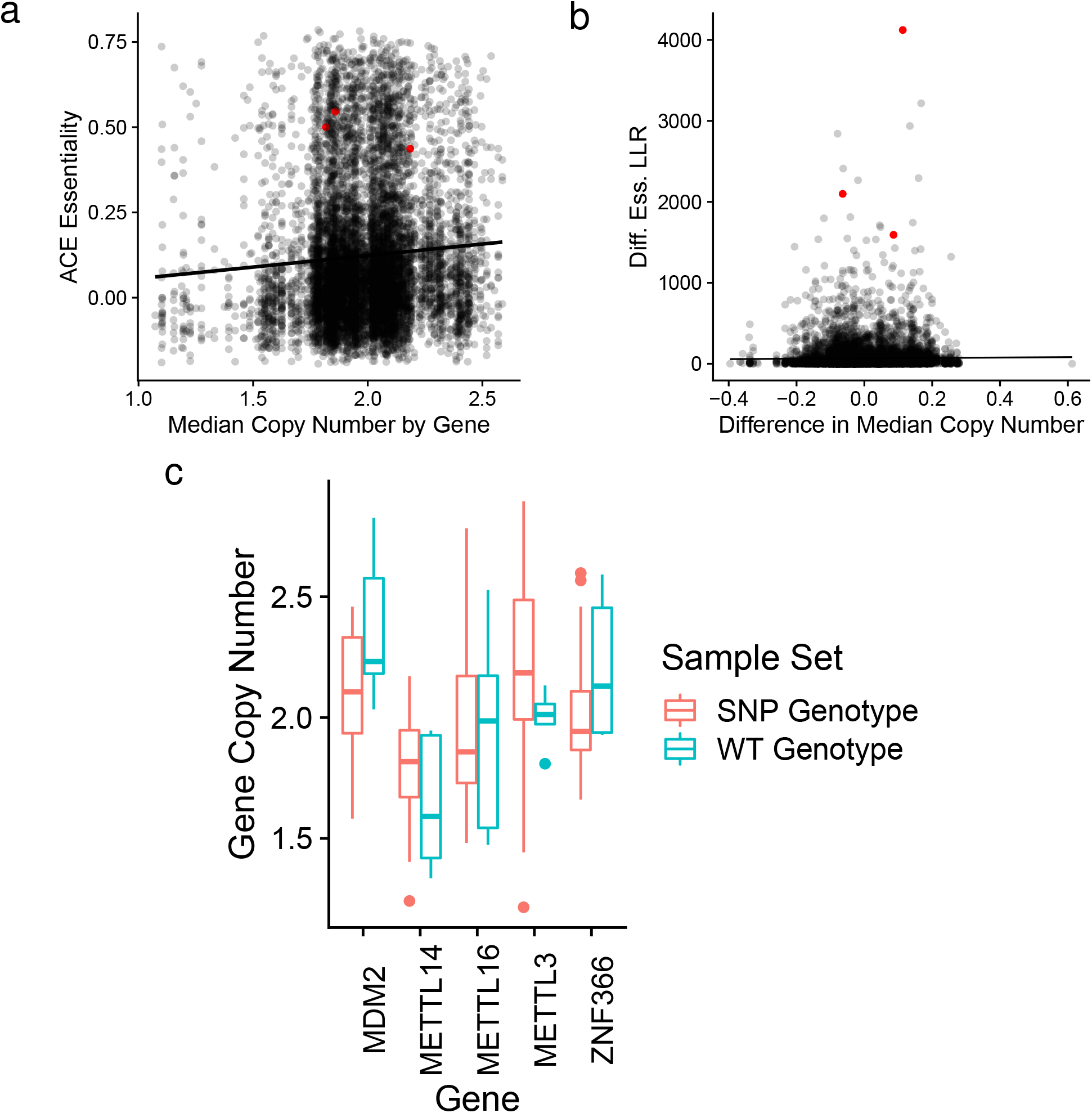
ACE Essentiality Estimates Avoid Copy Number Sensitivity by Pooling Across Sample Panels. (a) The ACE estimated essentiality of each gene (y-axis), with the median copy number of that gene across all NSCLC Adenocarcinoma samples with a mutation in *TP53* used to infer the essentiality value (x-axis). (b) The differential essentiality test statistic of each gene between wildtype and Δ*TP53* cell lines, determined by ACE. On the x-axis is the difference in median copy number between the compared panels of cell lines. In red are *METTL3, METTL14* and *METTL16*. The black line indicates the best linear relationship given a gaussian error model. (c) Copy number variation across each sample panel for the three candidate differentially essential methyltransferases, with *MDM2* as a positive control, and *ZNF366* as a negative control for differential essentiality. The range in copy number largely overlaps for all genes in these sample panels.

**Supplementary Figure S9.**
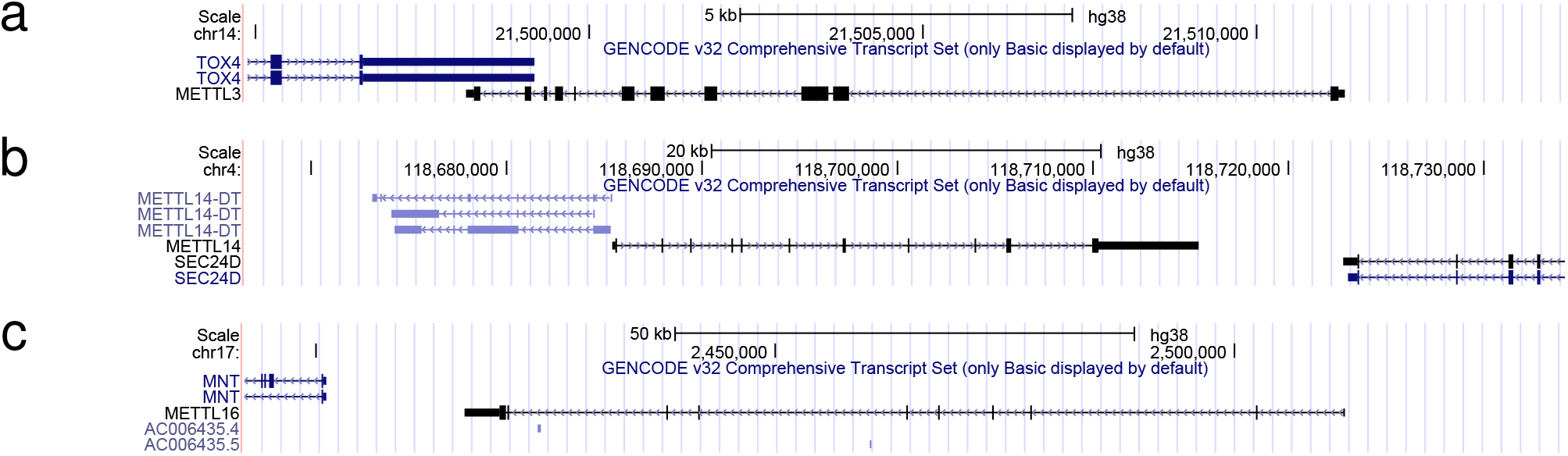
Differentially Essential Gene Candidates Independent of Flanking Genes. The majority of METTL3 (a), and all of METTL14 (b), and METTL16 (c) do not overlap with other coding genes. Image from the UCSC genome browser.

